# autoMEA: Machine learning-based burst detection for multi-electrode array datasets

**DOI:** 10.1101/2024.05.08.593078

**Authors:** Vinicius Hernandes, Anouk M. Heuvelmans, Valentina Gualtieri, Dimphna H. Meijer, Geeske M. van Woerden, Eliska Greplova

## Abstract

Neuronal activity in the highly organized networks of the central nervous system is the vital basis for various functional processes, such as perception, motor control, and cognition. Understanding interneuronal connectivity and how activity is regulated in the neuronal circuits is crucial for interpreting how the brain works. Multi-electrode arrays (MEAs) are particularly useful for studying the dynamics of neuronal network activity and their development as they allow for real-time, high-throughput measurements of neural activity. At present, the key challenge in the utilization of MEA data is the sheer complexity of the measured datasets. Available software offers semi-automated analysis for a fixed set of parameters that allow for the definition of spikes and bursts. However, this analysis remains time-consuming, user-biased, and limited by pre-defined parameters. Here, we present autoMEA, software for machine learning-based automated burst detection in MEA datasets. We exemplify autoMEA efficacy on neuronal network activity of primary hippocampal neurons from wild-type mice monitored using 24-well multiwell MEA plates. To validate and benchmark the software, we showcase its application using wild-type neuronal networks and two different neuronal networks modeling neurodevelopmental disorders to assess network phenotype detection. Detection of network characteristics typically reported in literature, such as synchronicity and rhythmicity, could be accurately detected compared to manual analysis using the autoMEA software. Additionally, autoMEA could detect reverberations, a more complex burst dynamic present in hippocampal cultures. Furthermore, autoMEA burst detection was sufficiently sensitive to detect changes in the synchronicity and rhythmicity of networks modeling neurodevelopmental disorders as well as detecting changes in their burst dynamics. Thus, we show that autoMEA reliably analyses neural networks measured with the multi-well MEA setup with the precision and accuracy compared to that of a human expert.

## 1 Introduction

In the human brain, highly orchestrated activity of neuronal networks lie at the basis of various functional neurological processes. In these networks, excitability is tightly regulated through a complex interplay between glutamatergic, excitatory neurons, and GABAergic, inhibitory neurons [1]. In the search to better understand processes contributing to a balanced network, multi-electrode arrays (MEAs) have provided a valuable tool to study the activity of neuronal networks as a whole [2, 3]. MEA devices allow for non-invasive measurement of electrical activity in neuronal cultures *in vitro* [4, 5]. Importantly, it allows one to follow the development of network activity as the culture matures and record responses of the network to compounds of interest [6, 7]. MEA has furthermore proven to be a valuable tool to model neurological diseases *in vitro* [8]. Many neurological diseases have been studied using neuronal networks derived from rodent brain tissue, such as Alzheimer’s disease, epilepsy and various neurodevelopmental disorders (NDDs) [8]. Since researchers are able to develop human induced pluripotent stem cell models of neurological diseases through differentiation into neuronal networks, MEAs have gained even more interest.

MEA electrodes record fluctuations in the electric field around them. When neurons on top of an electrode fire action potentials, the fast-flux of sodium and potassium ions across the membrane generates a typical change in the extracellular potential surrounding the MEA electrode, which is classified as a spike [8, 5, 9]. In typical MEA analysis software, spikes can be detected using a threshold for a minimal amplitude deviation from baseline noise (typically +/- 5 standard deviations) [5]. While frequencies of spiking activity can give information about the excitability of a network, this parameter is prone to fluctuations by technical and batch-to-batch variation [10]. More interesting and robust outcome measures are parameters that describe the activity of the network as a whole. During the initial phase of network development *in vitro*, spikes can be detected mostly in a random sequence. However, as the network starts to mature, periods of high-frequency spike trains are interrupted by periods of quiescence [9]. These high-frequency spike trains are classified as bursts. Typically, these bursts are recorded synchronously by multiple electrodes across the culture indicating the formation of a functionally connected network, hence, these are called network bursts (NB). Many parameters can be extracted from this type of activity, such as the synchronicity of the network, the rhythmicity of network activity, and characteristics such as burst duration and composition.

One major challenge faced when analyzing MEA data is the definition of bursts, about which no consensus has been reached in the research field [8]. Generally, bursts are defined based on a MaxInterval method, which defines a maximum inter-spike interval that is used as a threshold to classify a sequence of spikes as a burst. More extensive methods also include a maximum interval between bursts, the minimum duration of a burst, and a minimum number of spikes fired within a burst [11]. These thresholds can be chosen by an experimenter, or determined using adaptive burst detection algorithms [6, 12]. Electrophysiological mechanisms underlying burst dynamics are depending on multiple characteristics, such as neuronal excitability, synaptic transmission, and network connectivity. Hence, burst dynamics may differ between different types of neuronal cultures and change throughout network development [13]. For example, a study by Charlesworth and colleagues (2015) identified a unique feature of hippocampal neuronal cultures when compared to cortical cultures. From 11 days *in vitro* (DIV), hippocampal bursting dynamics were characterized by a theta rhythm, in which a single burst can be divided into multiple reverberations, i.e. short sequences of high-frequency spiking activity that closely follow each other, clustering into a burst [13]. While these reverberations can be detected using the same MaxInterval method, there is a higher chance of interference by spiking noise, e.g. single spikes occurring in between two reverberations thereby merging them together, because the inter-spike intervals will remain below the threshold. It is thus challenging to define a single set of parameters that can reliably define bursts over different experiments and culture types, and more complex burst dynamics may require more adaptive detection methods.

Currently, analysis is often carried out in software provided with the hardware (e.g. Multiwell Screen by Multi Channel Systems). In this software, parameters are set by the experimenter, based on visual inspection of the data, searching for the most ideal parameters for a certain dataset or by thresholding using the log inter-spike interval (ISI) [11, 6, 12]. However, this default set of parameters is often error-prone, especially when the data contains more complex bursting dynamics such as the reverberations in hippocampal cultures. For example, reverberations can also be detected using the MaxInterval method, but may be merged due to a single spike fired in between two reverberations. Current detection methods can only ignore such spiking noise if settings are manually altered by the experimenter through visual inspection. The visual inspection of the data to find the ideal set of parameters and adjust within a parameter’s defined range if necessary, is a very labor-intensive process, requiring file-by-file analysis of the data, and creates a risk for experimenter bias and reduces the objectivity of the analysis method. Therefore, often a default set is chosen taking for granted that multiple reverberations may be merged together in more noisy recordings.

Over the past several years, several analysis packages for MEA data have been published [14, 15, 16, 17]. While these packages provide more extensive and automated analysis options than software provided with the recording system, the MaxInterval and logISI burst detection methods integrated into this software generally use parameters that are not suitable for the detection of reverberations within bursts (e.g. interburst interval of 100ms). These methods are thus limited to simpler bursting dynamics, and not that of for example hippocampal bursts. Additionally, recent studies have started to identify reverberations in disease models of iPSC-derived neurons, which may further implicate the usefulness of a detection model that can accurately detect these more complex burst dynamics [18, 19, 20].

Machine learning (ML) models have become common in various domains, demonstrating remarkable efficacy and facilitating practical applications in everyday life. In scientific tasks, ML has spread through nearly every field, offering a valuable tool, particularly for tasks requiring automation, such as fine-tuning intricate devices [21, 22] or analyzing complex datasets [23, 24]. In connection to the problem of burst detection described above, ML emerges as a promising solution. Particularly, in scenarios employing the MaxInterval method, human experts must iteratively select parameters and inspect data quality until convergence to an optimal parameter set is achieved. One can exploit the optimal parameter determination process by selecting a set of MEA-data and correspondent optimal MaxInterval parameters and use it to effectively train a ML model to replicate the decision-making of human experts in parameter selection. Here, we have developed an automated analysis software tool, including optimized burst detection using a machine learning approach. We generated a sophisticated noise-resilient algorithm that takes a MEA signal or a spike train as input, and outputs the MaxInterval parameters that return bursts that would have been manually predicted by a human expert. This process is fully automated and does not need any manual assistance by a human operator. We present our algorithmic solution to the burst determination challenge and provide its implementation as a ready-to-use open-source package, autoMEA [25, 26], that the neuronal network community can immediately use and expand upon. We demonstrate that our approach works for a range of different input data (raw measurements averaged in different ways as well as binary spikes data) and we validate the model using existing datasets of two neurodevelopmental disorders (NDDs) across different time points in neuronal network development.

## 2 Results

### 2.1 Machine learning models

The autoMEA software detects bursts using two different methods: 1) the default method, with which detection is done using the same MaxInterval parameters as in the manual analysis software, and 2) burst detection based on MaxInterval parameters predicted by a machine learning model. In this work 3 different models were generated and implemented in the software. The three models are all built upon 1D-convolutional artificial neural networks, and have a different architecture depending on the input data: spikes30 model uses a 5-second binary spike trace averaged by 30 time-steps; signal30 uses real-valued signal averaged by 30 time-steps; and signal100 the real-valued signal averaged by 100 time-steps. Schematic depiction of our workflow is shown in Figure 2. The detailed information on the architecture and training of neural network machine learning models spikes30, signal30, and signal 100 is available in the Supplemental Information. The averaging of the original 5-second data was performed to reduce input size, thereby significantly reducing the computational power needed by the models. We tested different averaging window sizes to make sure the model’s performance was not compromised. All models’ output consist of a three-dimensional array, corresponding to 3 out of the 5 MaxInterval parameters. These predicted parameters are then applied in the standard MaxInterval method to detect reverberating bursts.

**Figure 1:**
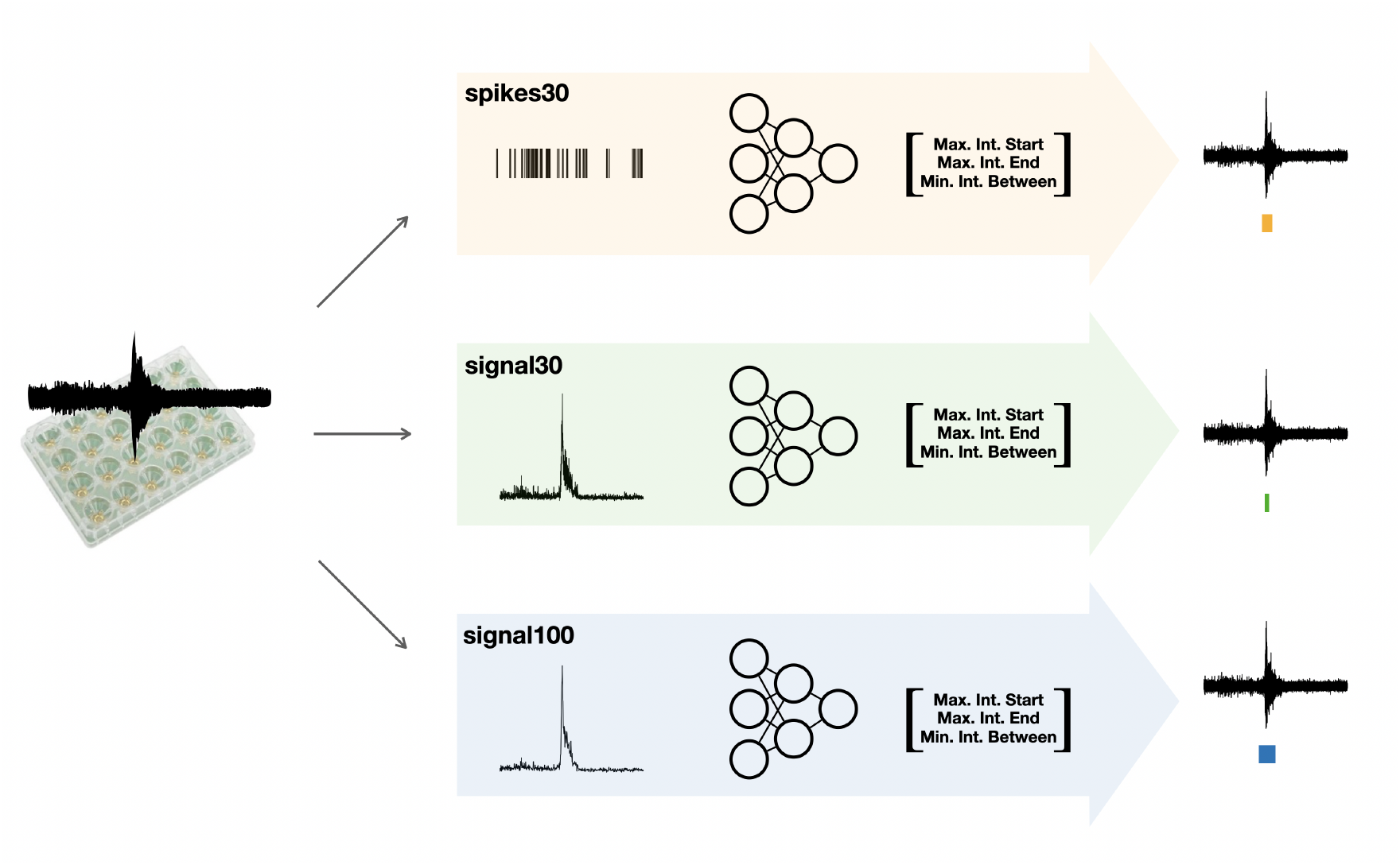
Schematic depiction of our workflow. After collecting the raw MEA data, we feed them into three different workflows: we post-process the data either into the form of spikes or averaged signal (over 30 or 100 time bins). We follow by training the neural network specific to each of the inputs (these models are referred to as spikes30, signal30, and signal100) to output key parameters for the MaxInterval method: maximum interval to start and end the burst and minimal time between the bursts. These parameters predicted by each machine learning model are then used for MaxInterval method that predicts the bursts.

**Figure 2:**
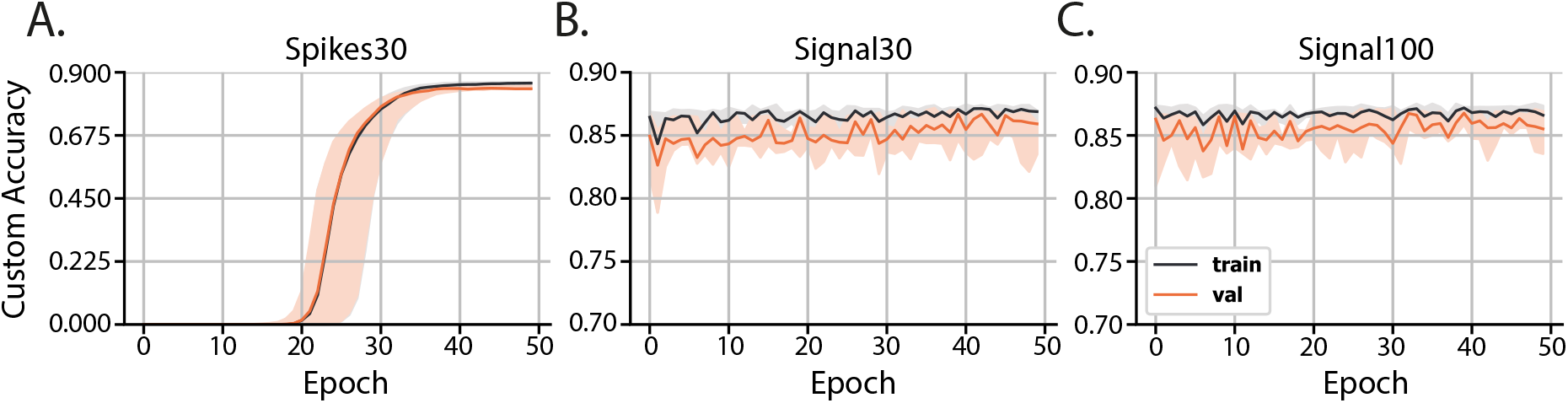
Custom accuracy for three machine learning models: spikes30, signal 30, and signal100. The solid line is the median of all custom accuracy values calculated for each epoch, and the shaded area is the range between the minimum and maximum values of custom accuracy for each epoch.

To train the machine learning models, a relatively small dataset comprising 797 burst samples was utilized. The dataset was built by using a functionality of our package that plots windows containing signal, spikes, and bursts detected using specific MaxInterval parameters. This feature was used to perform a standard post-processing analysis, where optimal MaxInterval parameters for detecting bursts were selected by the experimenter, and windows of 5-second duration containing signal/spikes/bursts and their corresponding parameters were saved for each sample.

The training of all three types of artificial neural network models (spikes30, signal30, and signal100) employed Mean Squared Error (MSE) as a loss function to measure the distance between the optimal human-selected parameters and those predicted by the network. A thorough hyperparameter tuning process was conducted by experimenting with different layers sizes and activation functions, followed by selecting the best overall hyperparameters for each model. To evaluate the efficacy of the predicted parameters in producing optimal bursts, simply relying on the loss function showing differences between sets of MaxInterval parameters was insufficient since there are multiple MaxInterval method parameter combinations that yield low loss and none of these combinations are captured by a single label consisting of experimenter’s parameter choice. Hence, a custom accuracy metric was introduced: it compares the bursts obtained obtained from parameters chosen by the artificial neural network and those selected by the experimenter. The learning curves showing the custom accuracy, for the training and validation set, are shown in Figure 2. From the learning curves, we observed that the spikes30 model gradually learned to predict *MaxInterval* parameters throughout the training process. The custom accuracy stayed very close to zero for the first 15 to 25 epochs, and then increased until converging to a value close to 0.86 (Figure 2A). In contrast, the signal30 and signal100 models, which use normalized signal as input, achieved a high value of custom accuracy (approximately 0.86) already in the first learning epoch, and the learning process was mostly visible by a shortening of the shaded area (statistical variation of network’s predictions), signaling that the accuracy of the signal30 and signal100 models was converging to a common value (Figure 2B-C).

#### Assessment of burst detection quality

Next, we assessed the accuracy of burst detection by the machine learning model in comparison to using the default MaxInterval parameter. We compared the default parameters and machine learning model as follows: an experienced experimenter was presented with one burst, the detection of which was presented in two different ways: using the default parameters, and the machine learning model’s predicted parameters. Being blind to the detection type, the experimenter then scored the detection as equal or gave a preference for one over the other detected burst. The quantitative overview of this comparison is shown in Figure 3. The experimenter scored 120 bursts for each of the three neural network models: spikes30, signal30, and signal100. The majority of bursts were detected equally well by the default method and either of the model’s detection as judged by the experimenter. For spikes30 model, there was an equal amount of the bursts that were better detected by the default and machine learning methods. For signal30 and signal100 models, the machine-learning-based parameter prediction slow slight statistical advantage. Overall, however, the detection by default method and by the machine learning models are comparable.

**Figure 3:**
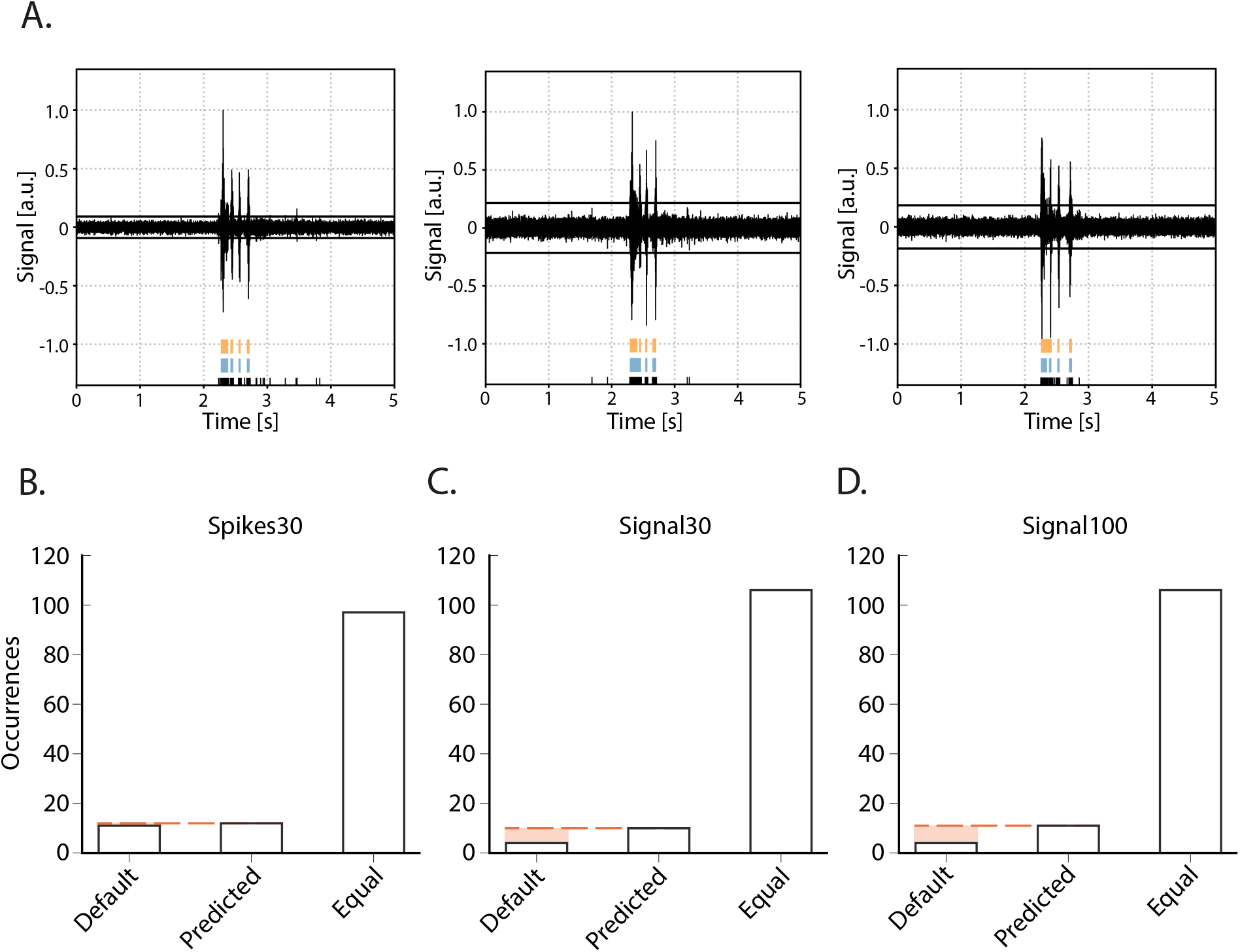
Burst quality metric for the parameter prediction models. A) 3 examples of bursts as presented for scoring to an experimenter. Raw signal of a network burst with detected spikes indicated in black at the bottom and above the reverberations as detected by either the default method or one of the detection models (spikes30, signal30, signal100). During scoring, the experimenter was blind to which color represented the default and spikes30/signal30/signal100 detection, and color could switch with each presentation of a new burst. e.g. A (left) default in blue, spikes30 in orange, (middle) default in blue, signal100 in orange, (right) default in orange, signal100 in blue. B-D) Burst quality score with the number of bursts that were scored as preferred with default method, with a predicted model, or equal for each method.)

### 2.2 Validation of parameter detection

#### Accuracy of spike and network dynamics detection by the autoMEA software

Network bursts are typical electrical activity patterns characterized by high-frequency spiking activity. Similar to the MCS Software, in the autoMEA software, spikes are used to detect bursting activity in the network. Thus, for the model to accurately detect network bursts, it was first of all important that the spikes could be correctly detected with the reproduction of the signal and threshold settings in autoMEA software. To this extent, we correlated the MFR of all wells in our hippocampal datasets, detected by manual analysis using the MSC software, to the MFR detected by the autoMEA software. We found a near-perfect correlation between the MFR detected by the manual analysis and autoMEA software (r(79) = 0.9939, p < 0.0001, Figure 4).

**Figure 4:**
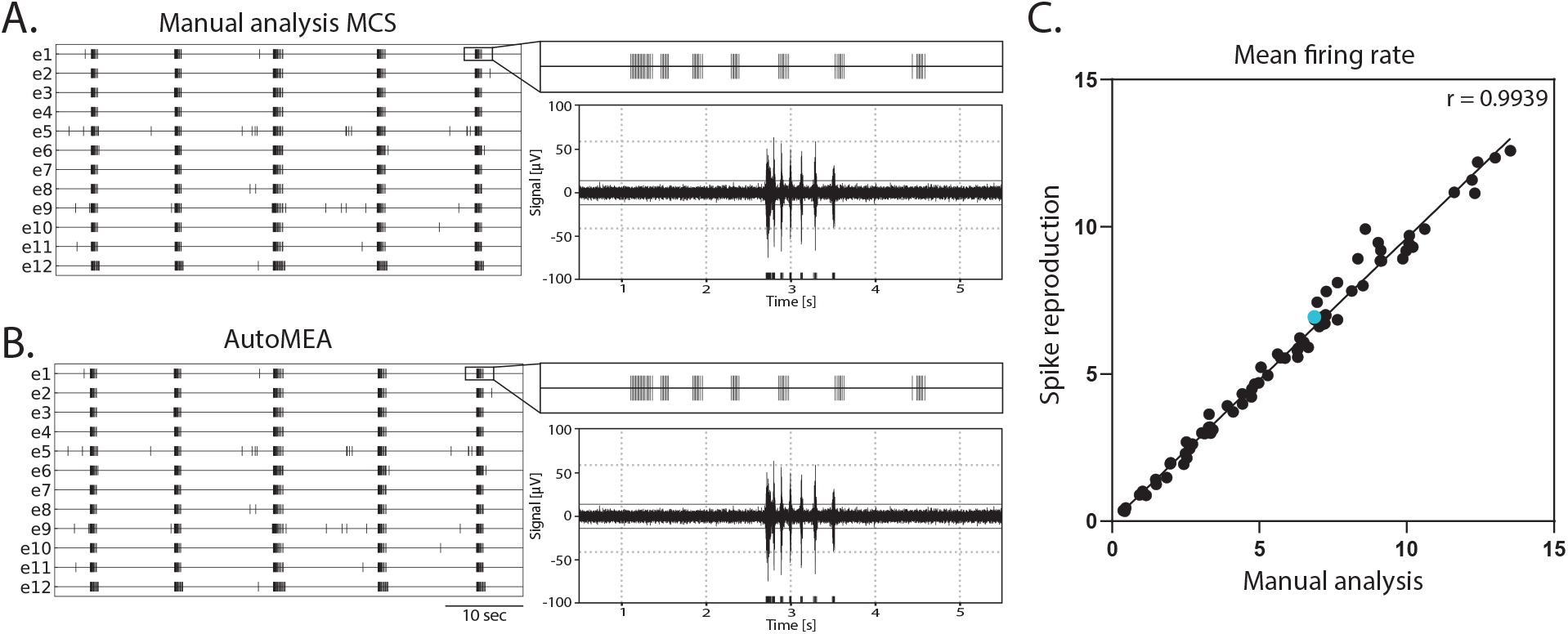
Correlation of the mean firing rate between spikes detected with manual analysis in MCS and autoMEA software. A) example raster plot (left) of spikes detected by MCS analysis with a zoom-in of the raster for a 5-second section above a 5-second section of the raw data (right). B) example raster plot (left) of spikes detected by the autoMEA software with a zoom-in of the raster for a 5-second section above a 5-second section of the raw data. C) Mean firing rate in Hz, detected by manual analysis in MCS software on the x-axis, and autoMEA software on the y-axis. The correlation between the two detection methods is near-perfect. The blue dot is the datapoint presented in figures A and B. N = 81 wells

Subsequently, we assessed the correlation between the manual analysis and the different detection methods: using default parameters, and either of the machine learning prediction models, for a set of outcome measures that can be used to describe neuronal network dynamics (Figure 5). Firstly, the % of random spikes (%RS, i.e. spikes not being part of a burst) and the network burst rate (NBR) can together describe the level of synchronicity in the network (Figure 5B1-2). We found that the detection of these parameters by any of the autoMEA models strongly correlated with the manual analysis. For both parameters, all autoMEA models showed correlations of r > 0.9 (statistics are presented in Table 1). All autoMEA models slightly overestimated the %RS, while on average fewer network bursts were detected (Figure 5B1-2).

**Table 1:** Relation of input, output and name of every model developed in this work.

| Approach | Input (length) | Output (length) | Name |
| --- | --- | --- | --- |
| Parameter Prediction | Binary spike array (1667) | Float parameter array (3) | spikes30P |
|  | Float signal array (1667) |  | signal30P |
|  | Float signal array (500) |  | signal100P |

**Figure 5:**
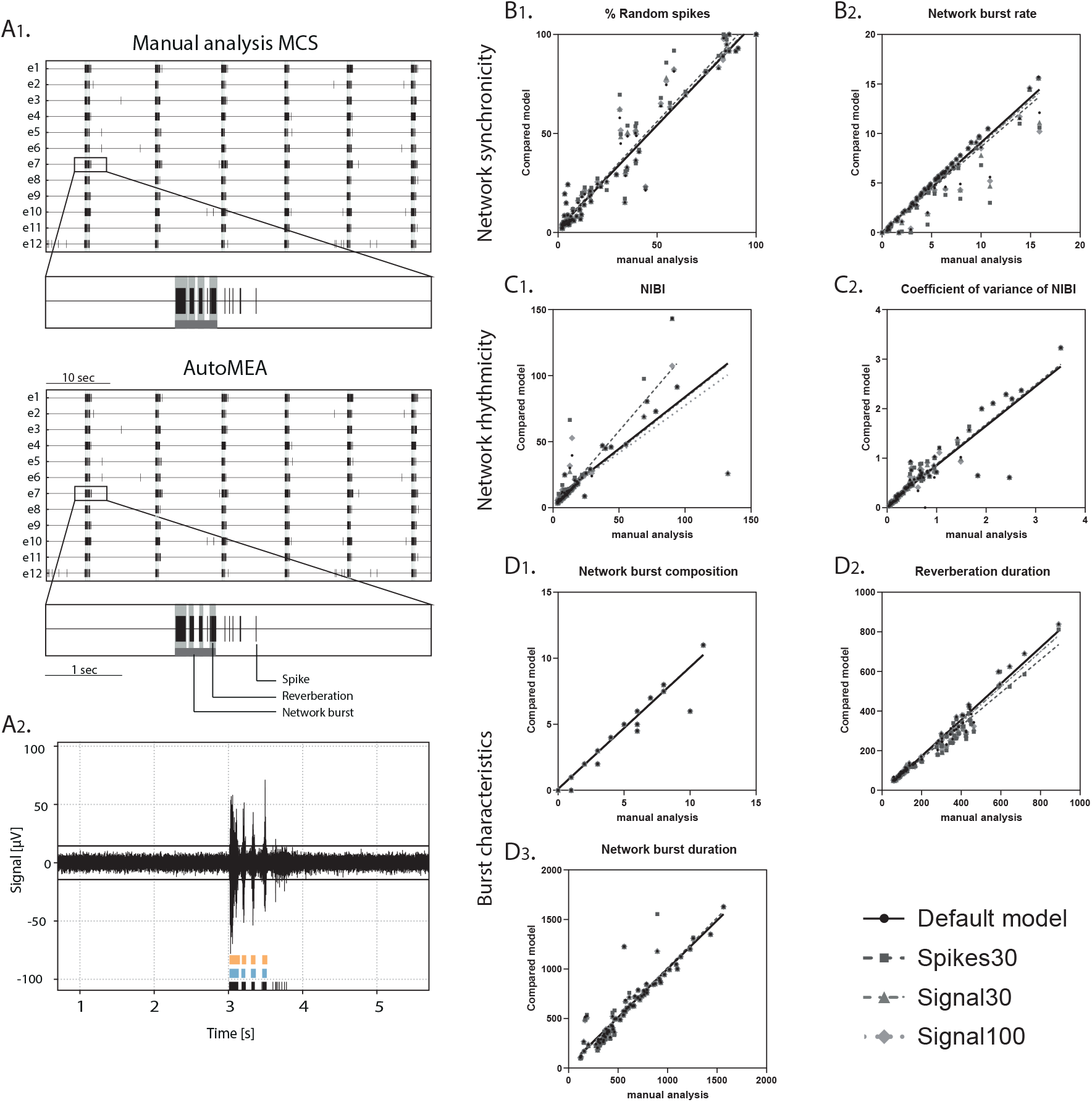
Validation of accurate parameter detection by the autoMEA models: A) 1. examples of raster plots and 5-second sections of a single electrode as detected by the Manual analysis in MCS (top) or autoMEA software (bottom). Black lines represent spikes, light gray bars overlaying raster plot represent reverberations, dark grey bar at the bottom of the zoom section represents the network burst. 2. raw data example of a network burst with reverberation detection presented at the bottom of the graph by manual analysis MCS in orange (top) and autoMEA analysis in blue (middle) and spikes in black (bottom). B) correlation between outcome parameters for network synchronicity 1. %RS, 2. Network burst rate. C) correlation between outcome parameters for network rhythmicity 1. NIBI, 2. Coefficient of variance of NIBI. D) correlation between outcome parameters for burst characteristics 1. network burst composition, 2. reverberation duration, 3. network burst duration. N = 81 wells.

| Method | %RS | NBR | NIBI | CoV of NIBI | NBD | NBC | Reverberation duration |
| --- | --- | --- | --- | --- | --- | --- | --- |
| Default | 0.9315 | 0.9720 | 0.9279 | 0.9239 | 0.9470 | 0.9911 | 0.9882 |
| spikes30 | 0.9331 | 0.9284 | 0.8864 | 0.9653 | 0.9446 | 0.0955 | 0.9842 |
| signal30 | 0.9333 | 0.9586 | 0.9228 | 0.9706 | 0.9499 | 0.9912 | 0.9880 |
| signal100 | 0.9340 | 0.9623 | 0.9184 | 0.9668 | 0.9512 | 0.9912 | 0.9867 |

Secondly, the rhythmicity of network burst firing can be described by the network inter-burst intervals (NIBI) and more specifically, the coefficient of variance (CoV-NIBI) thereof. These two parameters, determined by all autoMEA models, also strongly correlated with the results of the manual analysis (Figure 5C1-2, statistics are presented in Table 1). Interestingly, the outcome parameter NIBI showed a difference in the directionality of the change for the different models. The spikes30 model detected a slightly increased NIBI and showed the least strong correlation to the manual analysis (r(74) = 0.8864, p>0.0001). The default, signal30 and signal100 results showed strong correlations with manual analysis (r > 0.9). For these approaches, the detection resulted in a reduced NIBI compared to the manual analysis. Finally, network bursts can be characterized by their duration and reverberations. We again correlated the outcome of each autoMEA model to the manual analysis and found a strong correlation between the Network burst duration (NBD), Network burst composition (NBC i.e. reverberations / network burst), and reverberation duration (Figure 5D1-3, statistics are presented in Table 1). Here, both reverberation duration and NBC were lower when detected using any autoMEA model, while NBD was slightly higher.

Importantly, we observed that the difference between the prediction models and the manual analysis was mostly driven by the data processing method, as we observed that the default method already introduced small differences in burst detection compared to the manual analysis. The deviation between the default method and the prediction models was very small, showing the autoMEA models accurately reproduced the detection of reverberating bursts compared to detection by a default parameter set. Only for the outcome parameter NIBI did the spikes30 model deviate more from the default method and the signal prediction models.

Taken together, these results show that we can accurately detect reverberating bursts using the autoMEA software, in a way that is at least as good as a default set chosen by an experimenter through extensive visual inspection.

### 2.3 Validation of phenotype detection

#### Detection of neuronal network phenotypes in genetic models of NDDs

To validate the sensitivity of the autoMEA software for phenotype detection in disease models, we compared the analysis of the autoMEA software with manual analysis performed by an experienced researcher. Two different MEA datasets of NDDs were used for this comparison: 1) The RHEB-p.P37L model [27, 28], representing a disorder associated with severe refractory epilepsy due to hyperactivity of the mTOR pathway. The RHEB-p.P37L model has been well characterized on the MEA and showed increased spike- and bursting activity, premature synchronization of network activity, and loss of the reverberating burst pattern [29]. 2) The CAMK2g-p.R292P mouse model [30, 31] for a neurodevelopmental disorder associated with severe intellectual disability, autism, and general developmental delay. This model has not previously been characterized using multi-electrode arrays.

#### RHEB-p.P37L

In this validation experiment, we used a dataset of neuronal network activity recordings of the RHEB-p.P37L model at two recording days (days *in vitro*: DIV): DIV7 and DIV14. The hippocampal cultures were transduced at DIV1 with a virus inducing the expression of the patient-identified pathogenic RHEB-p.P37L variant [29], or a control virus.

First, we compared the network activity of control and RHEB-p.P37L neuronal networks, with bursts detected using the manual analysis to all autoMEA models at DIV14, a time-point at which control hippocampal cultures show reverberating bursts in a synchronous and rhythmic pattern. The averages of both genotypes were very similar, comparing the different autoMEA approaches to the manual analysis (Figure 6, for all outcome parameters, see Supplementary Figure 1). While there are slight differences in the exact numbers detected by the autoMEA software when compared to the manual analysis, these differences have the same directionality for both genotypes tested, e.g. the average reverberation duration slightly reduces for both the control and RHEB-p.P37L group. Furthermore, performing statistics on the difference between the two groups revealed that all methods accurately detected previously identified phenotypes: a decrease in NBC (Figure 6B) and an increase in reverberation duration (Figure 6C). Parameters that did not manifest a phenotype through manual analysis, similarly remained non-significant when analyzed using the autoMEA models (Supplementary Figure 1).

**Figure 6:**
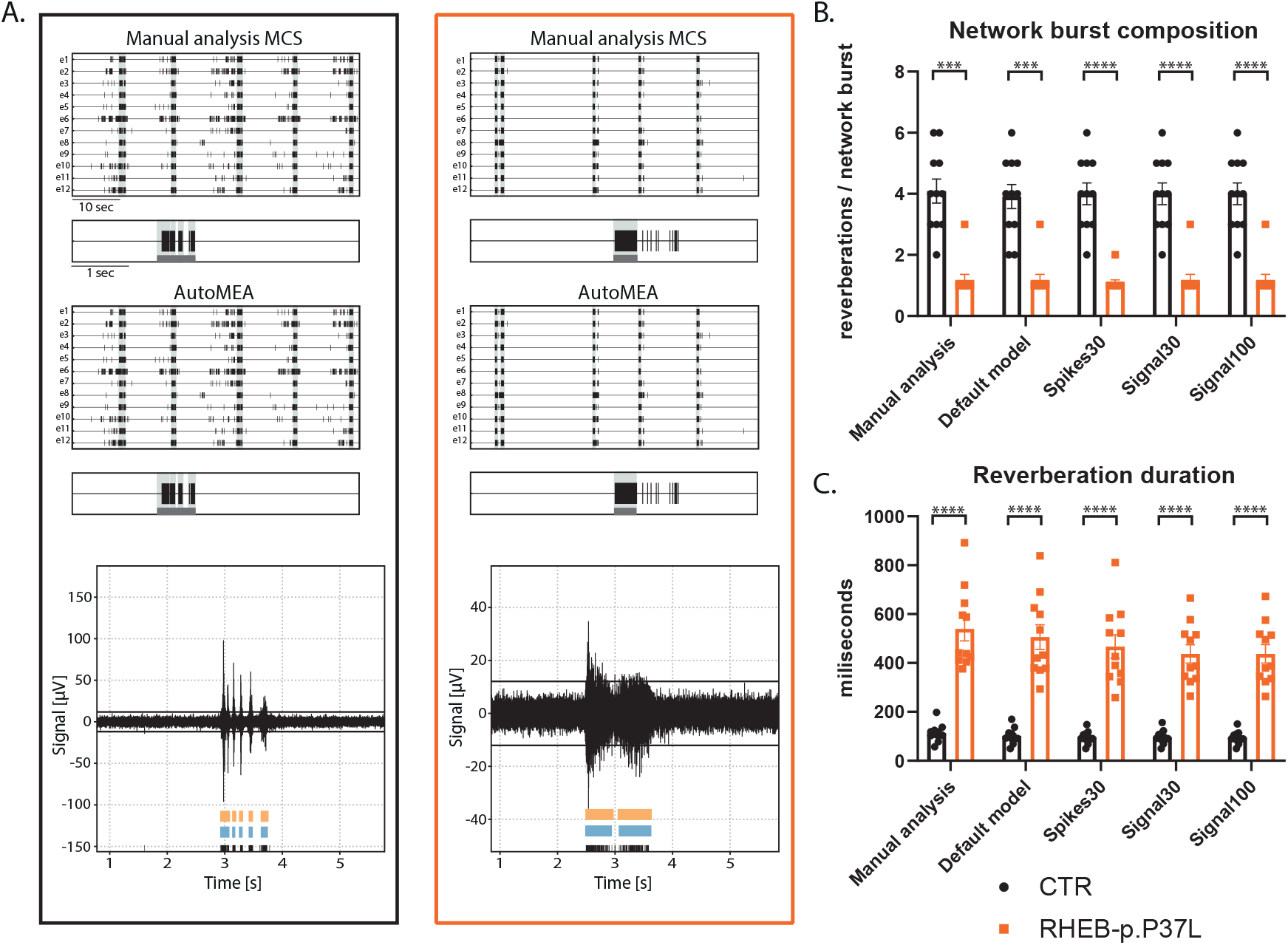
Validation of the detection of epilepsy-related phenotypes in a DIV14 set of the RHEB-p.P37L NDD model by the autoMEA software: A) Example raster plots with a 5-second section of a single electrode of a control (black, left) and RHEB-p.P37L (orange, right) well from manual MCS analysis and one autoMEA model, and a raw data trace example of a burst for both genotypes at the bottom, black lines at the bottom represent spikes, orange bars (top) represent reverberations as detected with manual MCS analysis and the blue bars (middle) reverberations detected using one of the autoMEA models. B) Comparison of the network burst composition for control and RHEB-p.P37L cultures detected using all different burst detection methods. C) Comparison of the reverberation duration for control and RHEB-p.P37L cultures detected using all different burst detection methods. N = 11 wells/group. Student’s t-test: ***p<0.0001, ****p<0.00001

To assess whether the software can also accurately detect bursts across the development of the culture, we included the analysis of the RHEB-p.P37L dataset at DIV7. At this time point, hippocampal cultures have not yet developed the reverberating network bursts and show more random spiking activity [13]. Similar to DIV14, we observed genotype averages very similar to the manual analysis for each parameter, and again, the directionality of change was the same for both genotypes (Figure 7, for all outcome parameters, see Supplementary Figure 2). Notably, also bursts in younger cultures, without reverberations appear to be accurately detected by the autoMEA models, which we trained to detect reverberating bursts. Furthermore, statistical comparison of the groups showed that phenotypes were accurately detected in DIV7 cultures (Figure 7, for all outcome parameters, see Supplementary Figure 2).

**Figure 7:**
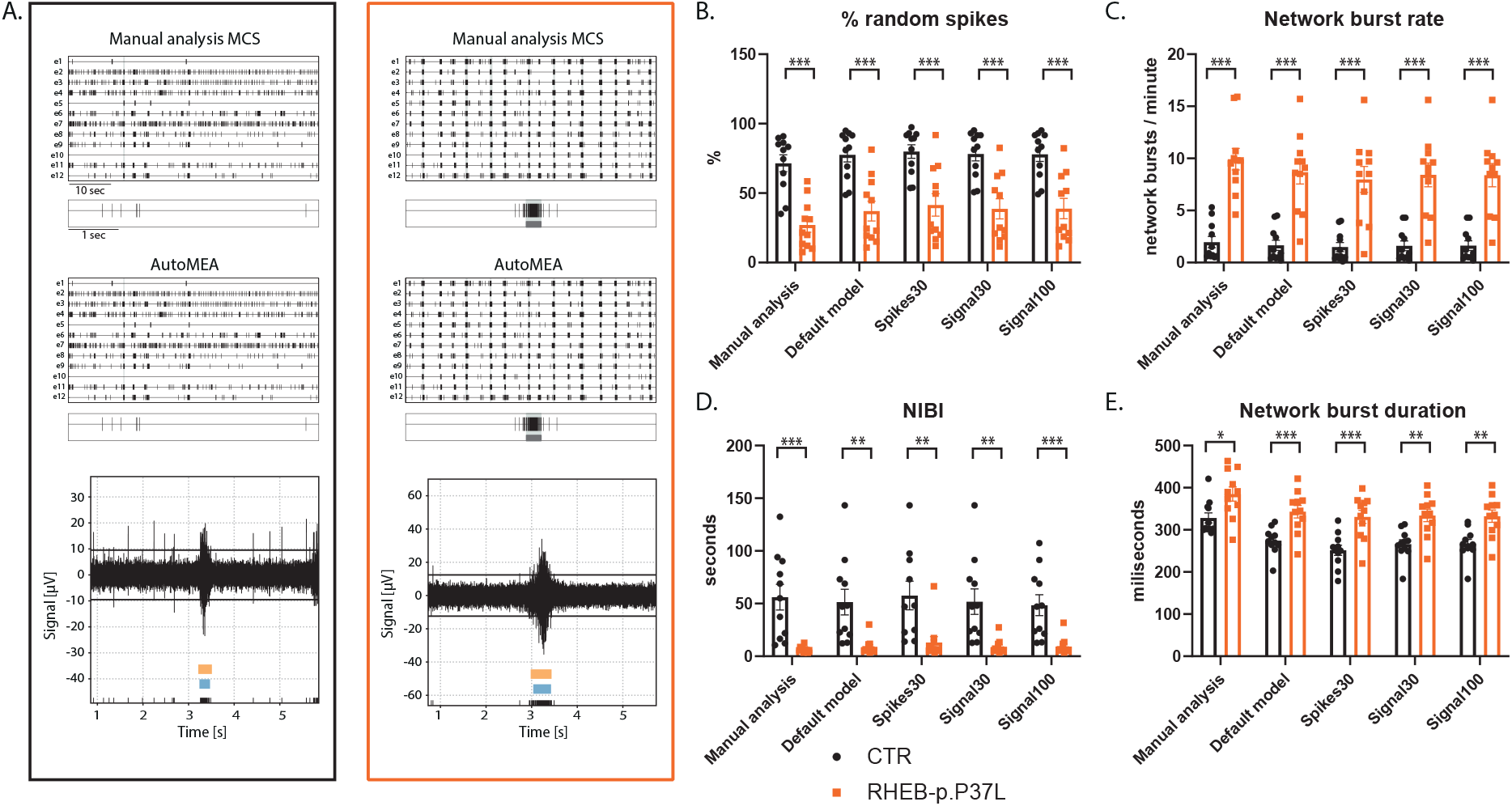
Validation of the detection of epilepsy-related phenotypes in a DIV7 set of the RHEB-p.P37L NDD model by the autoMEA software: A) Example raster plots with a 5-second section of a single electrode of control (black, left) and RHEB-p.P37L (orange, right) well from manual MCS analysis and one autoMEA model, and a raw data trace example of a burst for both genotypes at the bottom, black lines at the bottom represent spikes, orange bars (top) represent reverberations as detected with manual MCS analysis and the blue bars (middle) reverberations detected using one of the autoMEA models. B) Comparison of the %RS for control and RHEB-p.P37L cultures detected using all different burst detection methods. C) Comparison of the network burst rate for control and RHEB-p.P37L cultures detected using all different burst detection methods. D) Comparison of the NIBI for control and RHEB-p.P37L cultures detected using all different burst detection methods. E) Comparison of the network burst duration for control and RHEB-p.P37L cultures detected using all different burst detection methods. N = 11 wells/group. Student’s t-test: *p<0.05, **p<0.01, ***p<0.001 ***p<0.0001, ****p<0.00001

#### CAMK2g-p.R292P

We included a second NDD model, that has not yet been extensively characterized using MEA. Patients with this mutation suffer from severe intellectual disability (ID), autism spectrum disorder (ASD), and general developmental delay [30, 31]. In this second validation experiment, we used a set of neuronal network activity recordings at DIV18, from primary hippocampal neuronal networks transduced at DIV1 with a virus expressing either CAMK2G wildtype (CAMK2G-WT), the previously published pathogenic variant of CAMK2G, CAMK2G-p.R292P [31] or a control virus. Manual analysis of this novel dataset presented multiple phenotypes. We observed a decrease in the firing rate in both CAMK2G-WT and CAMK2G-p.R292P cultures compared to cultures transduced with a control virus (Supplementary Figure 3A). Furthermore, the expression of CAMK2G-p.R292P reduced network synchronicity as there was an increased percentage of random spikes (Supplementary Figure 3B-C). Interestingly, we identified decreased rhythmicity of network bursts for CAMK2G-WT, but not CAMK2G-p.R292P cultures, as shown by the significant increase in the CoV-NIBI (Figure 8B). We further observed apparent phenotypes in the network burst characteristics, namely that the NBD significantly decreased in CAMK2G-p.R292P cultures, while the reverberation duration increased (Figure 8C,E). CAMK2G-WT cultures displayed the opposite effect, an increased NBD while reverberation duration significantly decreased. Finally, we observed an increase in the NBC in the CAMK2G-WT cultures while this was drastically decreased in the CAMK2G-p.R292P cultures (Figure 8D). Also in this disease model, genotype averages were comparable between the manual analysis and the different autoMEA models and we could detect the same phenotypes in a novel dataset using the autoMEA models compared to the manual analysis.

**Figure 8:**
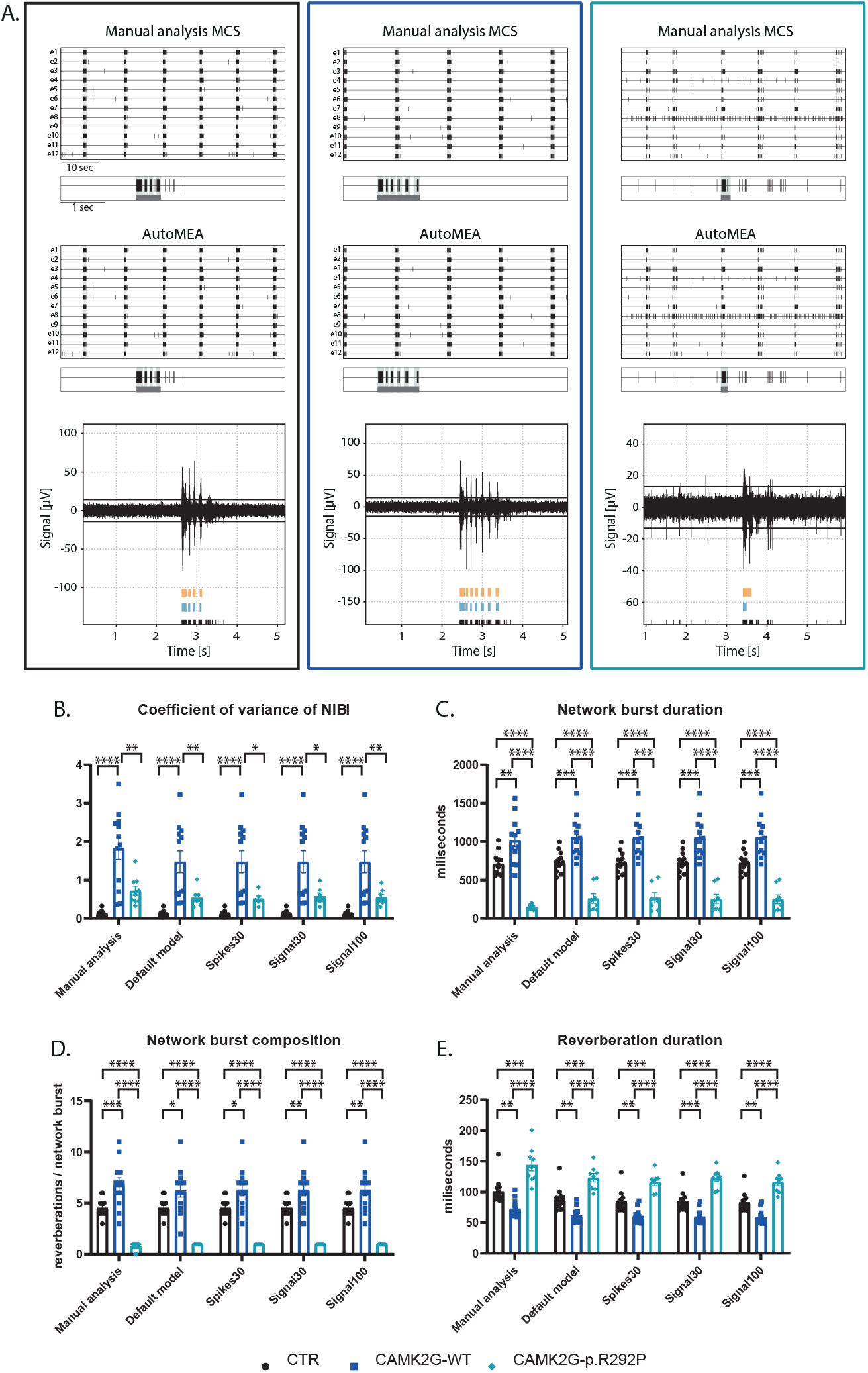
Validation of the detection of phenotypes in a DIV18 set of the CAMK2G-p.R292P NDD model by the autoMEA software: A) Example raster plots with a 5-second section of a single electrode of a control (black, left) and CAMK2G-WT (dark blue, middle), and CAMK2G-p.R292P (light blue, right) well from manual MCS analysis and one autoMEA model, and a raw data trace example of a burst for all genotypes at the bottom, black lines at the bottom represent spikes, orange bars (top) represent reverberations as detected with manual MCS analysis and the blue bars (middle) reverberations detected using one of the autoMEA models. B) Comparison of the %RS for control, CAMK2G-WT and CAMK2G-p.R292P cultures detected using all different burst detection methods. C) Comparison of the network burst rate for control, CAMK2G-WT and CAMK2G-p.R292P cultures detected using all different burst detection methods. D) Comparison of the NIBI for control, CAMK2G-WT and CAMK2G-p.R292P cultures detected using all different burst detection methods. E) Comparison of the network burst duration for control, CAMK2G-WT and CAMK2G-p.R292P cultures detected using all different burst detection methods. N(control) = 13 wells, N(CAMK2G-WT) = 12, N(CAMK2G-p.R292P) = 12. One way ANOVA: *p<0.05, **p<0.01, ***p<0.001 ***p<0.0001, ****p<0.00001 in the autoMEA software in the detection of NDD-related phenotypes.

In summary, the autoMEA model accurately detects phenotypes in hippocampal cultures of two different NDD models. Importantly, it detects phenotypes at multiple time points in the development of the cultures. This data shows that the autoMEA software is a reliable tool to analyze hippocampal MEA datasets. We did not observe striking differences between the performance of the different prediction models incorporated

#### Cortical data

To investigate if the performance of the model is specific to the hippocampal burst dynamics of the dataset that was used to generate the model, or if it can accurately detect bursts across datasets with different burst dynamics, we included a set of recordings from a cortical dataset at DIV14. While hippocampal cultures generate spontaneous reverberating bursts, cortical cultures do not present this reverberating pattern (Figure 9). A cortical dataset of 18 wells was analyzed using the manual settings used to analyze the hippocampal data and analyzed using the autoMEA models. On top of that, the MaxInterval method parameters in the manual detection analysis were adjusted to more accurately detect bursts with cortical burst dynamics. To this extent, the maximum interspike intervals to start and end a burst were set to 100 ms, and the minimum interval between bursts was set to 200 ms, from here referred to as ISI100 analysis.

**Figure 9:**
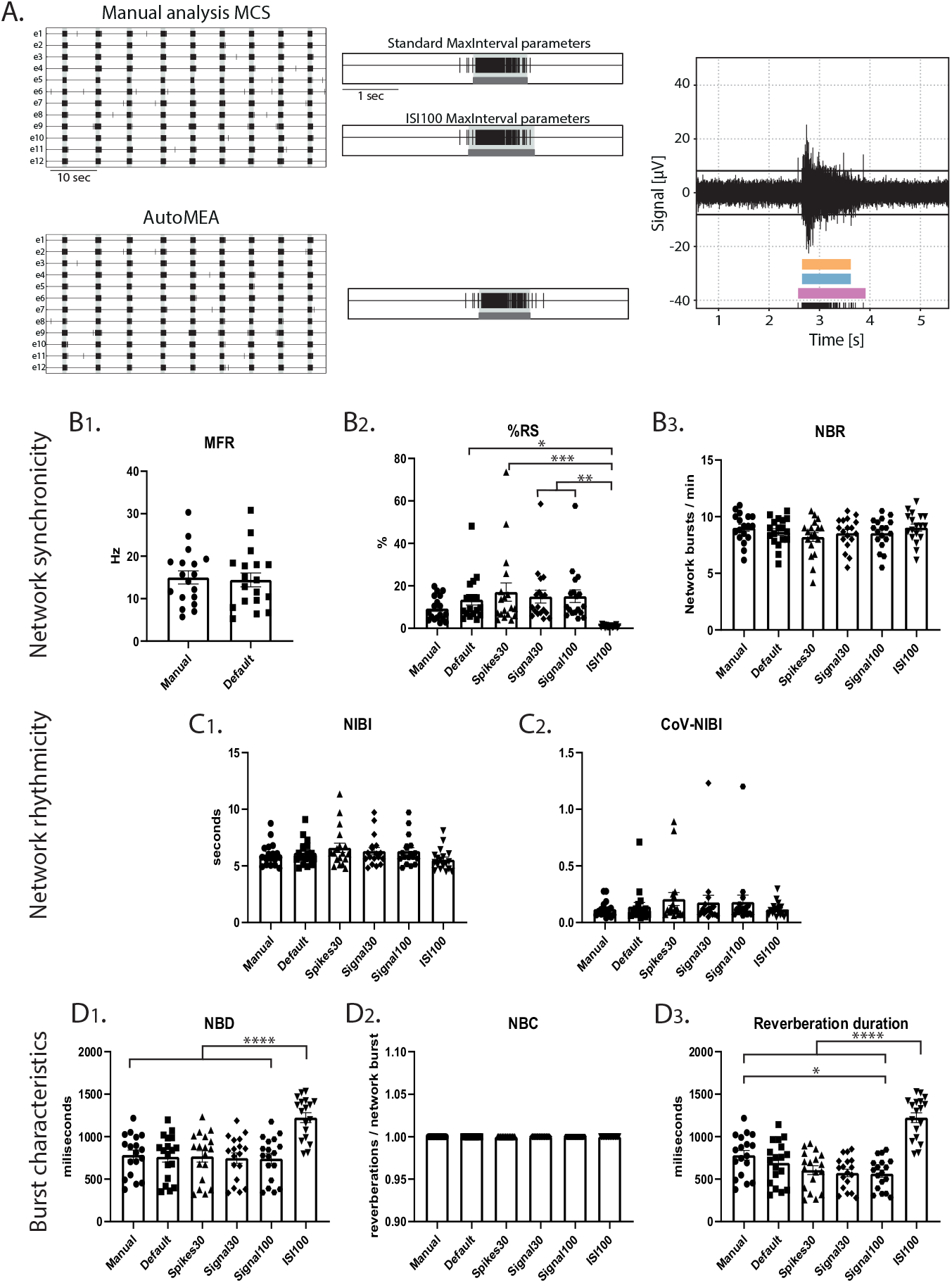
Testing the performance of autoMEA burst detection on a cortical dataset. A) Example raster plots with a 5-second section of a single electrode, analyzed using manual MCS analysis and one autoMEA model, and a raw data trace example of a burst for all genotypes at the bottom, black lines at the bottom represent spikes, orange bars (top) represent reverberations as detected with manual MCS analysis, the blue bars (middle) reverberations detected using one of the autoMEA models and the pink (bottom) the manual detection in MSC using ISI 100. B) Comparison of the MFR, %RS, and NBR using all different burst detection methods. C) Comparison of the NIBI and CoV-NIBI using all different burst detection methods. D) Comparison of the NBD and NBC and reverberation duration using all different burst detection methods. N = 18 wells. One way ANOVA: *p<0.05, **p<0.01, ***p<0.001 ***p<0.0001, ****p<0.00001

Similar to the hippocampal datasets, the detection of bursts with all its analyzed characteristics was very similar between the manual analysis and the autoMEA models for most of the wells. The adaptation of the MaxInterval parameters significantly affected the detection of the following parameters: as expected when the threshold for the ISI increases, network burst duration significantly increased, paired with a decrease in the % RS, as more spikes were identified as part of the burst (Figure 9). The other parameters regarding burst frequency and rhythmicity were not significantly affected by increasing the ISI threshold, and as such, the autoMEA model accurately detected these outcome parameters in our cortical cultures.

Interestingly, the dataset also presented a clear outlier (Supplementary figure 4), which came up most obviously in the read-out parameters %RS and CoV-NIBI. Inspection of this well showed that some network bursts were not detected using the detection models, and most network bursts were missed when the spikes30 model was used. The amplitude of the spikes within these bursts appeared lower, however this was not quantified.

In summary, our findings demonstrate that our automated quantitative machine-learning based analysis software tool, autoMEA, is a reliable tool to analyze MEA datasets in a high-throughput manner, presumably without inter-experimenter bias. Moreover, the model’s detection proficiency extends to effectively capture bursts with varying dynamics, showing its ability to generalize across different datasets.

## 3 Discussion

In this project, we generated automated detection software that can be used for the analysis of neuronal network activity recorded using a multiwell MEA system. More specifically, our focus was on creating a package with the capability to accurately identify intricate burst dynamics inherent to hippocampal neuronal networks. With this approach, we additionally aimed to reduce the manual aspect of the analysis and improve the efficiency with which the data can be analyzed. We showed that the different detection models included in our autoMEA software can accurately detect network burst activity from the MEA signal, comparable to a manual data analysis, using a defined set of MaxInterval parameters. The outcome from the autoMEA models showed very strong correlations with the manual analysis. Additional scoring of the model’s performances revealed that in most cases, the models could predict network bursts as well as the default MaxInterval parameters, in some cases performing even better than default as judged by an experienced experimenter. More importantly, the autoMEA models were able to identify the same phenotypes that were identified using manual analysis in two separate datasets for NDDs. Finally, the models could accurately detect bursts with different dynamics that appear throughout the maturation of a neuronal network, as was shown by the detection of bursts in a dataset of DIV7 neuronal networks.

MEA is a valuable tool for investigating neuronal network activity and is often used for disease modeling or toxicological assessments. However, burst detection remains a challenge in the field, as burst dynamics can vary depending on the type of cultures that are recorded [13, 32]. While previously, several MEA-analysis tools have been generated, they focused on the analysis and visualization of bursts with simpler network dynamics [14, 15, 16, 17]. In these models, bursts are detected based on the MaxInterval and/or logISI burst detection methods that have not been optimized for the detection of reverberations. Here, we presented a model that accurately detected reverberations based on the MaxInterval method but using adaptive parameters to optimize reverberation detection, a parameter that is sensitive to spiking noise.

With autoMEA software, we also provide experimenters with an open-source user-friendly software package. While the software that is provided with the commerical hardware outputs timestamp files that need to be post-processed to extract relevant parameters, our model does not require any need for coding expertise or manual data processing to post-process the output into quantifiable outcome parameters that are relevant to describe neuronal network dynamics. Besides outputting timestamp files, it automatically generates an additional output file with the network parameters describing network synchronicity, rhythmicity, and burst characteristics. Furthermore, one can input multiple recording files into the software, letting it analyze multiple recordings at the same time. Additionally, manual analysis requires the experimenter to actively adjust settings during the analysis of each file, while with autoMEA software, the analysis is performed completely autonomously without the experimenter’s input. Running the software may take from minutes up to a few hours, depending on the size of the dataset. Therefore autoMEA software enhances the throughput of MEA-data analysis. For a demonstration of the user-friendliness of autoMEA, see Supplemental Information.

Our MEA package is an open source package that can be completely adapted depending on the user’s needs. The key feature is that the machine learning models can be fine-tuned or retrained using new datasets. This may be preferred when datasets with different burst dynamics than the murine hippocampal cultures are analyzed. Researchers can build upon the current dataset, which could enhance the accuracy of the model to detect bursts with different burst dynamics, and importantly, can further reduce any experimenter bias as the model is now trained based on the analysis of an experienced researcher. By blinding the experimenter to bursts presented during the training, we tried to ensure the objectivity of the burst quality metric introduced in this study.

In this study, the machine learning model was generated to specifically detect the more complex burst dynamics observed in hippocampal neuronal networks, in which bursts generally consist of multiple reverberations. To broaden the usefulness of the autoMEA software, we showed that our package could also accurately detect many of the outcome parameters in a cortical dataset. However, as burst dynamics are different in cortical cultures, the MaxInterval parameter threshold for the ISI with which this type of data is analyzed is typically higher [6], which affected significantly the outcome parameters %RS and the duration of network bursts and reverberations. For different types of data, it may be necessary to retrain the models with a specific dataset, however, the autoMEA software could still be used without retraining if a predetermined set of MaxInterval parameters is known. In this case, the software can be run using the default method, similar as to what was done using the manual MCS analysis settings in the default method of the autoMEA software in this paper. We did not observe obvious differences between the performance of the different prediction models (spikes30, signal30, signal100). Both using binary spike input and real-valued signal, the machine learning approach was trained to accurately detect parameters describing network dynamics. The only small difference was identified in the detection of the NIBI, wherein the NIBI was increased using the spikes30 model compared to default, while it was reduced in the signal models. However, these deviations were small and did not affect the detection of phenotypes in our NDD models. Therefore, we consider all autoMEA models suitable for the analysis of MEA-data.

The autoMEA machine learning approach for detecting bursts from signal or spikes demonstrated robust generalization across diverse datasets. We introduced a customized accuracy metric that evaluates the difference between bursts detected by manually set MaxInterval Parameters and those predicted by our machine learning models, enabling precise performance assessment. Despite being trained on a modest dataset derived from a limited set of recordings, these models accurately replicate the experimenter’s MaxInterval Parameter selections for analysis. It’s noteworthy that all the machine learning models examined in this study are based on simple convolutional neural networks that are well established in the machine learning community and straightforward to implement. Despite their simplicity, all the machine learning models assessed in this study have shown impressive accuracy. Notably, the autoMEA package facilitates easy fine-tuning of the used machine learning models with additional data or their substitution with more advanced architectures, ensuring adaptability to evolving research needs.

With recent technological advancements that allow the development of neuronal networks derived from iPSCs, the MEA system has become more popular as a functional readout in disease modeling studies using stem cell methods [8]. Interestingly, in multiple disorders, the appearance of reverberations, otherwise referred to as fragmented bursts or super bursts, was identified as a phenotype [18, 33]. Therefore, we believe that our software can be of interest to a broader audience. The flexibility of our software allows users to use the models trained with the datasets presented in this paper, but also retrain it using their own dataset to optimize detection in datasets with different burst dynamics. Additionally, adding training data onto the current dataset may increase the ability of the software to analyze more complex or diverse datasets, and may result in better convergence of the model onto a dataset with varying burst dynamics. Thus, we provided here an effective software tool for multi-well MEA analysis that is user-friendly, high-throughput, and adaptable to the researcher’s preferences.

## 4 Methods

### 4.1 MEA data collection

#### MEA recordings of primary hippocampal neurons

Primary hippocampal and cortical neuronal cultures were prepared from embryonic day (E) 16.5 FvB/NHsD wild-type mice according to the procedure previously described [28] [34]. Neurons were plated in a multiwell multi-electrode array (MEA) plate with an epoxy base (Multichannel Systems, MCS GmbH, Reutlingen, Germany) in a density of 35,000 neurons / well. Cultures were maintained in neurobasal medium (NB, GIBCO) supplemented with 2% B27, 1% penicillin/streptomycin and 1% glutamax (NB+++) and placed in an incubator at 37 °C with 5% CO2.

Each MEA well is embedded with 12 PEDOT-coated gold electrodes of 100 *μ*m in diameter and 1 reference electrode. Recording electrodes are arranged in a 4×4 grid, spaced 700 *μ*m apart.

Twice weekly, neuronal network activity of the cultures was recorded after which one-third of the medium in each well was replaced with fresh NB+++.

MEA plates were recorded using the Multiwell-MEA headstage in a recording chamber at 37 C with 5% CO2. Recordings were started after 10 minutes of acclimatization in the MEA set-up. Neuronal activity was recorded for 10 minutes at a sampling rate of 10 kHz and the signal was filtered with a 4th order low-pass filter at 3.5kHz and 2nd order high-pass filter at 100 Hz [29].

#### Manual MEA data analysis using the MCS software

MEA data was manually analyzed using the MultiChannel System software package. Analysis was performed on the full 10-minute recording period. Baseline noise was calculated as the average of 2×200 ms segments without activity at the start of the analysis period, and a threshold of +/- 5 SD from baseline was used to detect spikes. Network reverberations were detected using the MaxInterval method that is incorporated into the MCS software. For the dataset used in this study, the most ideal parameters for reverberation detection were identified as:

Max. interval to start = 15 ms

Max. interval to end = 20 ms

Min. interval between = 25 ms

Min. duration = 20 ms

Min. number of spikes = 5

For the cortical dataset, manual analysis of the same wells was run with the settings adapted to Max. interval to start and end a burst = 100 ms and Min. interval between bursts at 200 ms.

Whenever at least two-thirds of the channels in a well were participating in synchronized activity, of which at least half of them were simultaneously active, it was classified as network activity.

Output from the MCS Software was then further processed using a custom-written script in MATLAB R2021a. Network reverberations were combined into network bursts when the interval to the next reverberation was <300 ms.

Using the custom-written processing script in MATLAB, multiple outcome parameters were extracted from the data that can describe the network development and dynamics in the culture. We calculated 8 different outcome parameters that could be classified into 3 categories: 1) Network synchronicity, described by the mean firing rate (MFR), network burst rate (NBR), and the % random spikes, i.e. spikes that are not part of a burst (%RS). 2) Network rhythmicity, described by the network interburst interval (NIBI) and more specifically the coefficient of variance thereof (CoV-NIBI). 3) Burst characteristics, to which the network burst duration (NBD), reverberation duration, and network burst composition (NBC: reverberations/network burst) are descriptive.

### 4.2 Machine-learning automation

The ultimate goal of MEA data analysis is to quickly and robustly detect bursts for large amounts of measured data. While the ideal MaxInterval parameters are hard to unify across the dataset, due to the spiking noise, it is still possible to adapt them manually to the different levels of noise. These manual adjustments can be automatized if one is able to develop an algorithmic mapping from MEA signal (or spikes) to the parameters during the processing of the dataset. Machine learning is a powerful tool that is able to approximate complex multivariable functions as well as to generalize well under the influence of noise. In our approach, we chose to use MEA data, such as processed signals and spikes, as input to a supervised neural network, trained to output optimal MaxInterval parameters for each data sample. This way, we allow for the parameters to be continuously adapted without the constant attention of a human operator. In this approach, the model developed closely mimics what an experimenter does when analyzing MEA data: it finds the parameters that help extract the best burst configuration from the dataset.

Below we describe how we generated a dataset to be used to train and test the models developed, and details about the methods implemented.

#### Dataset generation

In order to train and test the models in this work, we generated a dataset in which we selected five-second windows of MEA activity, during which bursts occur, and together with the experimenter expert, the values of the first three MaxInterval parameters (Max. interval to start, Max. interval to end, and Min. interval between bursts) are adjusted until an optimal burst detection set is obtained. This process is repeated multiple times, adding at each iteration one sample to the dataset - for this work, a total of 797 samples were generated, using recordings for different systems at different DIVs. For each selected window, all the data useful to train and evaluate the developed models is saved. For the sampling rate considered, 10kHz, a five-second window corresponds to 50 thousand timestamps, which is why most of the data is saved as arrays with lengths equal to 50 thousand. Specifically, MEA signal is saved a float array, while spikes and bursts as binary arrays, with an element equal to zero in case there is no spike/bursts activity occurring at the correspondent timestamp, and equal to one in case there is activity. Finally, the MaxInterval parameters are saved as an integer array with length equal to three - since we are just interested in the first three parameters. Afterwards, part of the original dataset was post-processed, to get variables in the form of input/output used by the different models developed, and divided into Training/Validation/Test sets. In details, the signal arrays were normalized between 0 and 1, and all temporal arrays (signal, spikes and bursts) were averaged by either 30 or 100 timestamps. In the end, we have three different inputs for each approach considered. In Table 1 we show the various inputs and outputs used for each model developed, assigning a name for each specific model.

#### Prediction of MaxInterval parameters

The models implemented consist of convolutional neural networks that receive as input MEA data, and map it into three of the five MaxInterval parameters. Different models were developed based on the different input choices, as described in the Dataset Generation section. All models were developed using the Tensorflow/Keras framework. The models were trained in a supervised learning regime, where a loss function - Mean Square Error in this case - is defined to calculate an error between the convolutional networks’ predicted output and the target output - the manually selected MaxInterval parameters. This error is then used to update the internal parameters of the model until the predicted output converges to the target. This iterative procedure used to optimize the model parameters is called training. During training, the model calculates different loss values for the samples taken from the Training Set and the Validation Set, with the main difference being that the model parameters are just updated based on the loss retrieved from the Training Set only. Moreover, the iterative training process is divided into epochs, where one epoch is an instance for which the model used the totality of the Training/Validation Set.

While designing a convolutional neural network many hyperparameters have to be defined, such as the architecture of the network, which optimizer is used to change the internal parameters, and how many samples are used before updating the parameters. In order to find the best hyperameter for each model implemented - depending on the input type, we used a package called hyperas, which works within the keras framework and makes it possible to define a set of values for each hyperparameter, and scan which combination of these values return the best models, based on the final loss value. Then, using hyperas, we trained each of the three models one hundred times using different hyperameters values, and post-selected three best cases for each one, based on the behavior of the training and validation loss.

The three best cases for each model were trained again, now calculating a new metric to characterize the accuracy of the model. Accuracy is a metric used to quantify the performance of a machine learning model, however, it is just meaningful when it is used for classification, when the output is a discrete value correspondent to a class, and the objective is to distinguish between different classes. In the Parameter Prediction approach, the models developed are performing what is called regression, when the output is a real value, used to estimate the value of a variable. However, the error that defines a distance between the predicted MaxInterval parameters and the target values is not enough by itself to define how the models are performing. The final goal of the automation developed in this work is to obtain optimal bursts, independently by the combination of MaxInterval parameters used to detect these bursts. We defined a custom accuracy metric, which compares the predicted bursts - in this case, the bursts detected using the predicted MaxInterval parameters, and the target bursts - the bursts detected using the target MaxInterval parameters. Considering the binary representation of the burst arrays, the custom accuracy is defined as

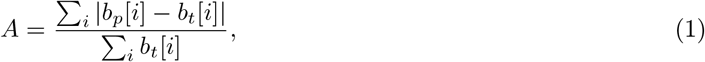

where *b*_*p*_[*i*] is the binary burst array, obtained using the predicted parameters, at index *i*, and *b*_*t*_[*i*] is the binary burst array obtained using the target parameters, at index *i*. This metric quantifies how many timestamps the bursting state differs between the bursts detected using predicted and target parameters, normalized by the number of timestamps in which the target bursts are active (equal to one). The normalization is necessary, given the sparse nature of the binary burst arrays, to avoid high accuracy values in cases where the predicted bursts are a full-zero array (no burst activity).

Calculation of the custom accuracy while training a model takes a considerable amount of time, since for each sample used a burst has to be detected and compared to the target one. That is why we just perform the custom accuracy calculation for the three best-performing cases for each model. Then, from the new training procedure, one best model for each input type is chosen, based on both the loss and the custom accuracy, and is trained five more times to obtain averaged values of loss/accuracy and test its consistency.

To further quantify the model performance, we defined a new metric, called Burst Quality, using the test set (never seen by the model), in which we detect bursts using both the MaxInterval parameters predicted by the ML-model, and those used as default by the experimenter expert. Both sets of bursts are shown to the expert together with the correspondent signal and spikes, and the expert votes on which burst detection better represents that in the specific time trace. For this we built a GUI that shows difference signals/spikes figures, randomly shuffling the position/color with which bursts detected using predicted/default parameters are plotted.

### 4.3 Model validation

To assess whether the model could accurately detect phenotypes in models for NDDs that were identified using the manual analysis in the MCS software, datasets of two different NDD models were used: RHEB-p.P37L and CAMK2G-p.R292P. The RHEB-p.P37L pathogenic variant has previously been identified in focal cortical dysplasia type 2 and is associated with severe epilepsy [27, 28], and has been extensively characterized using the Multi-electrode array [29]. The CAMK2G-p.R292P pathogenic variant has been identified in patients with severe intellectual disability [30, 31]. These disorders were modeled through a lentivirally induced expression of the RHEB-p.P37L, CAMK2G-WT or CAMK2-p.R292P genes, compared to transduction with a control virus. Recordings from different days *in vitro* were included in this study to verify the accurate detection of bursts throughout the development of the culture. Manual and autoMEA analysis were done using data from DIV7 and DIV14 for the RHEB-p.P37L model, and at DIV18 for the CAMK2G-p.R292P model. Additionally, the convergence of the model onto a cortical dataset was tested using a wild-type dataset of cortical data recorded at DIV14.

### 4.4 Statistics

Statistical analyses were performed using GraphPad Prism 5 (GraphPad Software, Inc., CA, USA). The normality of the data was assessed using the Shapiro-Wilk test. The correlation for each outcome parameter comparing the model analysis to the manual analysis was analyzed using Pearson’s r or the non-parametric alternative if the normality assumption was not met, and the linear relationship was plotted using simple linear regression. Statistical analysis of disease phenotypes was performed using a Student’s t-test (RHEB-p.P37L data), or One-way ANOVA (CAMK2g-p.R292P and cortical data). For all statistical analyses, alpha was set at 0.05. The specific tests used for each experiment are specified in the figure legends or the results section. Values are represented as averages ± SEM. Sample sizes for each experiment are indicated in the figure legends.

