## Supplementary material for "autoMEA: Machine learning-based burst detection for multi-electrode array datasets"

May 8, 2024

#### 1 Models' architecture and hyperparameters

From the hyperameter optimization step while training the machine learning models used in this work, it was possible to choose three best models for each input type, for which the training was repeated calculating the custom accuracy. Then, one single model is chosen for each input type, based on the highest value of custom accuracy achieved. An overview of the final models' hyperameters is shown in Table 1.

#### 2 autoMEA full analysis method

In this section, we showcase an example of how to use the autoMEA package in practice. The easiest way to use autoMEA is to perform a full analysis of a series of datasets and corresponding wells, using one of the machine learning models available with the package for burst detection.

The standard workflow consists of preparing a *csv* file containing datasets and corresponding wells to be analyzed. The structure to be followed is shown below in the example file called `datasets_and_wells.csv`.

Input file example

```
#dataset, wells
dataset_1.h5, A1 A2 B2
dataset_2.h5, B1 C2
```

This file is used to analyze the wells `A1`, `A2` and `B2` from `dataset_1.h5`, and wells `B1` and `C2` from `dataset_2.h5`.

Table 1: (DRAFT - have to recheck every hyperameter in the office pc) Hyperparameters for the best model for each input type. CL refers to the Convolutional Layer and FC to the Fully Connected Layer.

| | | Spike30 | Signal30 | Signal100 |
|---|---|---|---|---|
| CL1 | Filters | 128 | 128 | 128 |
| | Kernel size | 3 | 7 | 7 |
| CL2 | Filters | 64 | 64 | 64 |
| | Kernel size | 5 | 7 | 7 |
| | Max Pooling | Yes | No | No |
| | Dropout | 0.33 | 0.18 | 0.18 |
| CL3 | Filters | - | 16 | 16 |
| | Kernel size | - | 9 | 9 |
| FC1 | Units | 64 | 32 | 32 |
| | Dropout | 0.12 | 0.62 | 0.62 |
| Optimizer | | RMSProp | SGD | SGD |
| Learning Rate | | 1E-6 | 1E-4 | 1E-4 |
| Batch size | | 2 | 4 | 4 |
| Trainable parameters | | 6865731 | 921299 | 323795 |

If the *csv* file, the datasets to be analyzed, and the machine learning model saved as a *h5* file are present in the same folder, a script to run the analysis can be:

Python Code

```
import automea

# create automea analysis object
am = automea.Analysis()

# define machine learning model to use, and load it
am.model_name = ‘signal30.h5’
am.loadmodel()

# define which output will be saved
am.analysis_params[‘save_stats’] = True
am.analysis_params[‘save_net_bursts’] = True

# run analysis
am.analyze_dataset(‘datasets_and_wells.csv’)
```

The analysis is performed using the model called `signal30.h5`, and a statistics file and a network bursts file are produced as output, as specific by setting the correspondent items of the `analysis_params` attribute as `True`.

The network bursts are saved as *csv* file with information about the datasets and well analyzed, and information about each network burst. An example of the network bursts file structure is shown below, with ellipses used to omit less significant columns present in the file.

Network bursts output file example

```
Dataset, Well Label, Start time[µs], Duration[µs], Spike Count, ...
dataset_1.h5, A1, 981800, 395800, 498, ...
dataset_2.h5, C2, 465000, 120500, 112, ...
```

The statistics output file contains several higher-order statistical quantities that are calculated during the analysis. An example of the file structure is shown below, using again ellipses to omit some of the columns present in the file.

Statistics output file example

```
Dataset, Well Label, Firing Rate[Hz], Stray spikes[%], Number net bursts, ...
dataset_1.h5, A1, 7.8, 11.92, 49, ...
dataset_2.h5, B1, 8.2, 12.33, 45, ...
```

This example shows how easy and practical it is to use autoMEA as a ready-to-use analysis package. One of the main goals of the package is to place itself as an accessible alternative to researchers with limited coding skills. At the same time, more advanced functionalities can be accessed, and modified depending on users' needs. A series of tutorials showcasing how to use autoMEA to perform tailored analysis of MEA datasets can be found on the package website **READTHEDOCS_REFERENCE**.

### 3 Supplementary figures

Supplementary figure 1

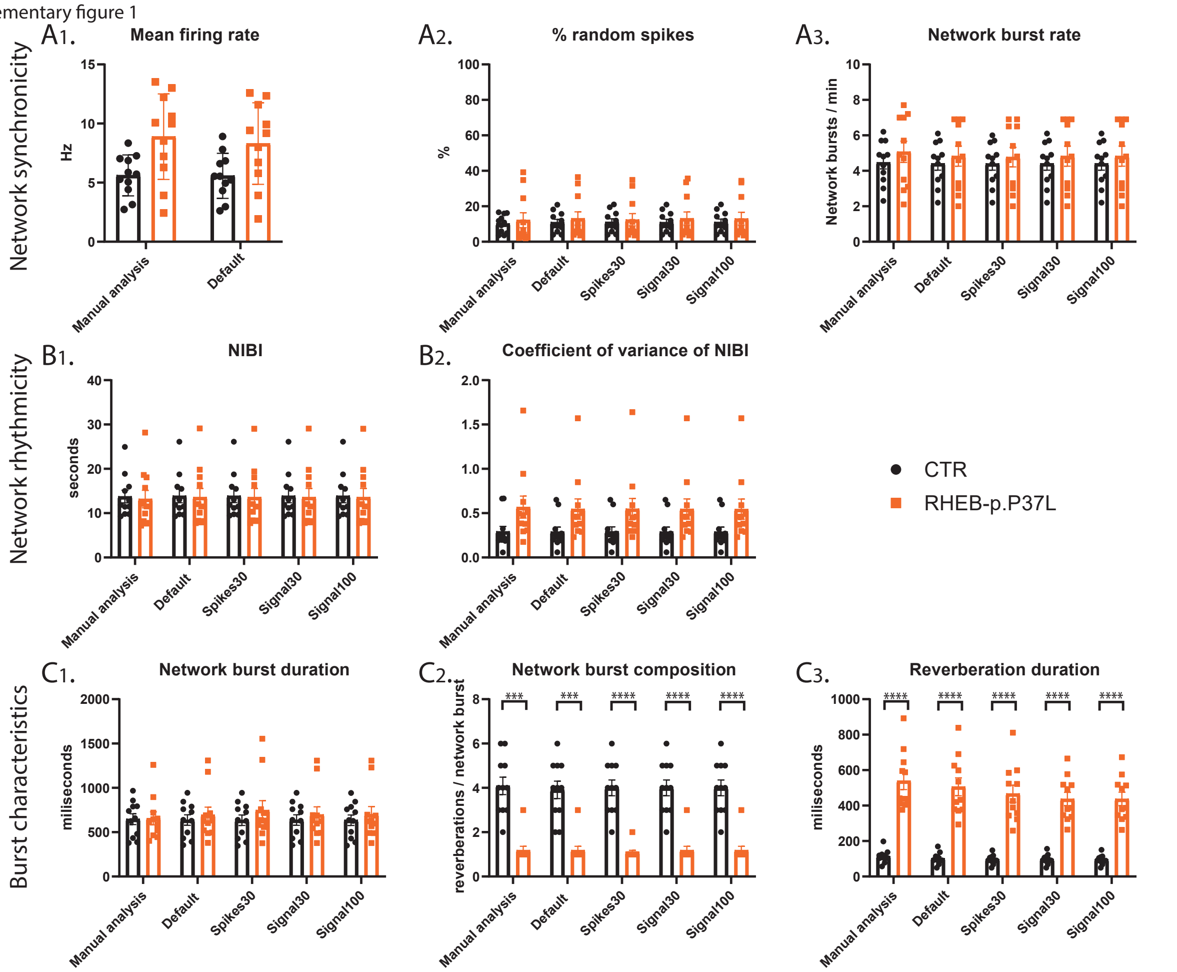


Figure 1: Validation of the detection of epilepsy-related phenotypes in a DIV14 set of the RHEB-p.P37L NDD model for all outcome parameters by the autoMEA software: A) detection by manual analysis and autoMEA for outcome parameters describing network synchronicity. B) detection by manual analysis and autoMEA for outcome parameters describing network rhythmicity. C) detection by manual analysis and autoMEA for outcome parameters describing burst characteristics. N = 11 wells/group. Student's t-test: *$p<0.05$, **$p<0.01$, ***$p<0.001$ ***$p<0.0001$, ****$p<0.00001$

Supplementary figure 2

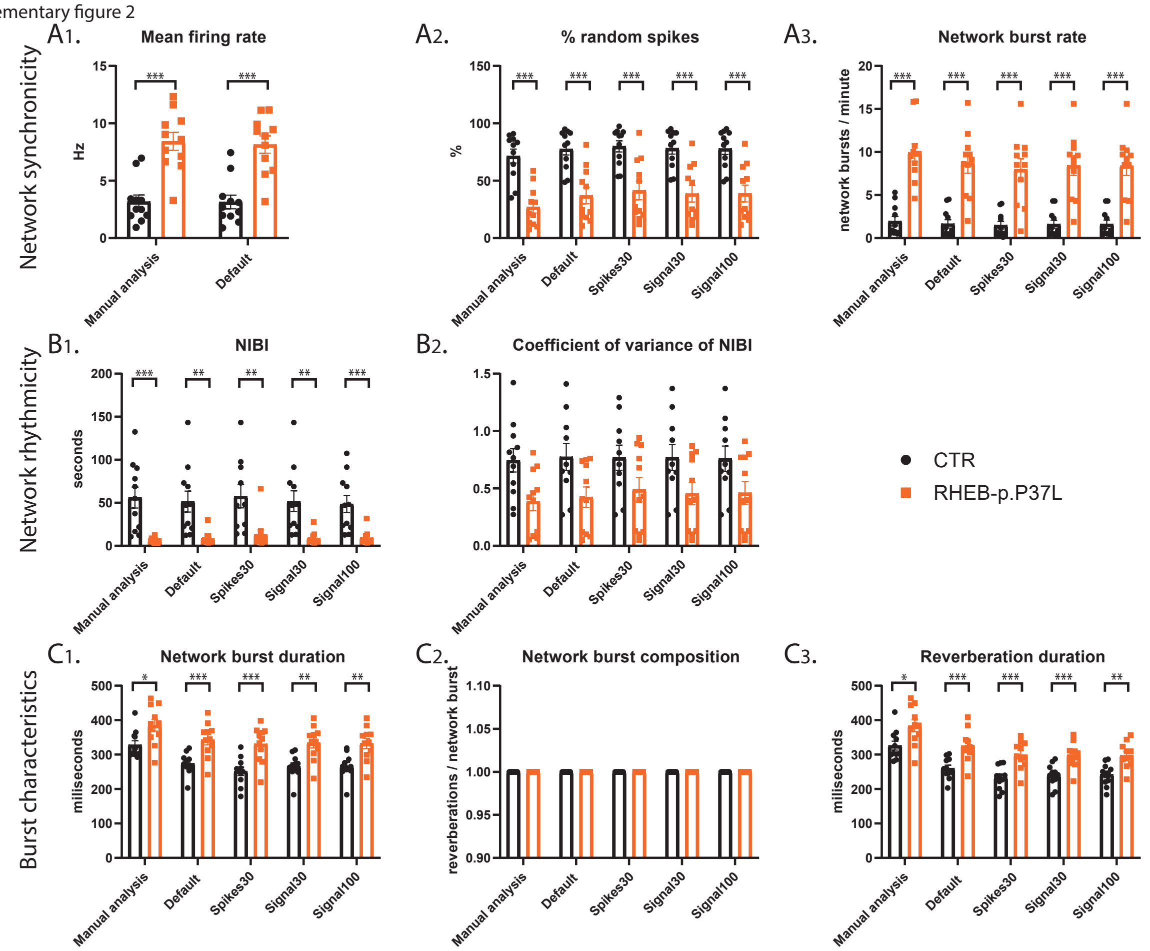


Figure 2: Validation of the detection of epilepsy-related phenotypes in a DIV7 set of the RHEB-p.P37L NDD model for all outcome parameters by the autoMEA software: A) detection by manual analysis and autoMEA for outcome parameters describing network synchronicity. B) detection by manual analysis and autoMEA for outcome parameters describing network rhythmicity. C) detection by manual analysis and autoMEA for outcome parameters describing burst characteristics. N = 11 wells/group. Student's t-test: *p<0.05, **p<0.01, ***p<0.001 ***p<0.0001, ****p<0.00001

Supplementary figure 3

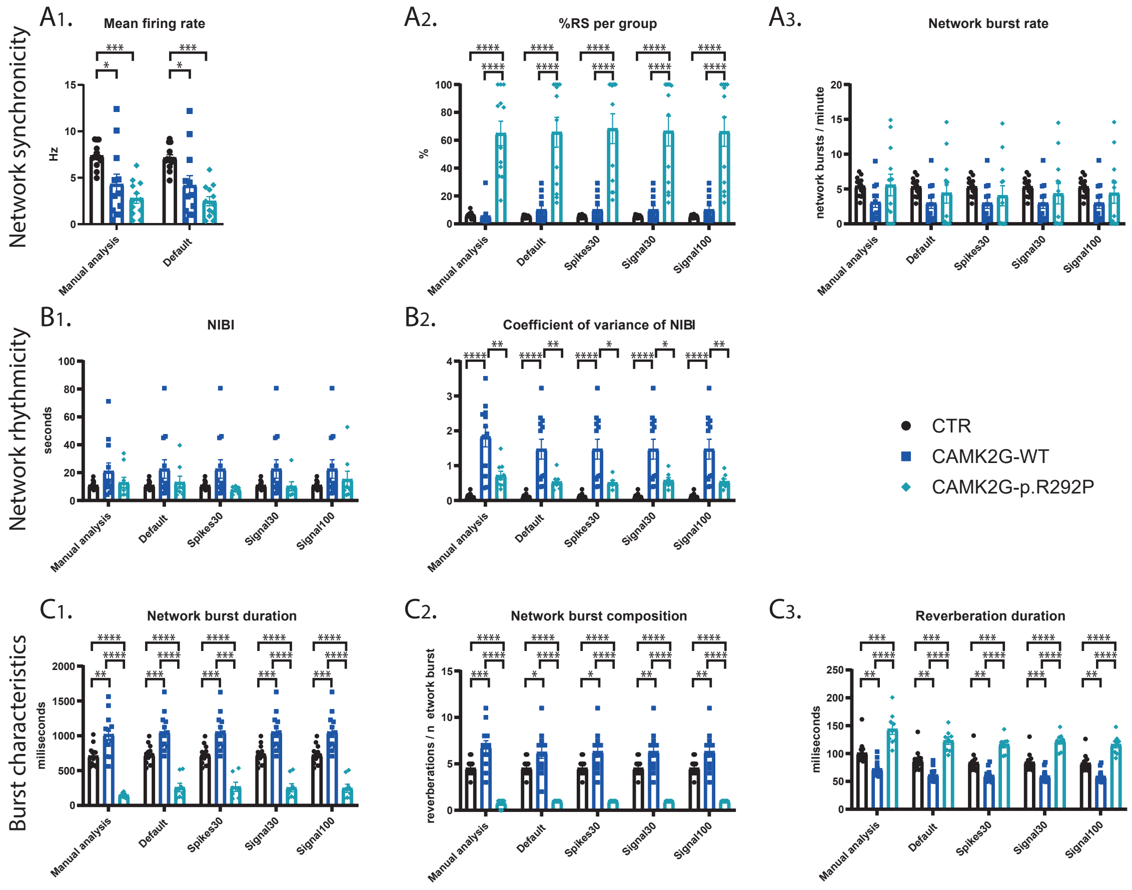


Figure 3: Validation of the detection of epilepsy-related phenotypes in a DIV18 set of the CAMK2G-p.R292P NDD model for all outcome parameters by the autoMEA software: A) detection by manual analysis and autoMEA for outcome parameters describing network synchronicity. B) detection by manual analysis and autoMEA for outcome parameters describing network rhythmicity. C) detection by manual analysis and autoMEA for outcome parameters describing burst characteristics. N(control) = 13 wells, N(CAMK2G-WT) = 12, N(CAMK2G-p.R292P) = 12. One way ANOVA: *p<0.05, **p<0.01, ***p<0.001 ***p<0.0001, ****p<0.00001

Supplementary figure 4

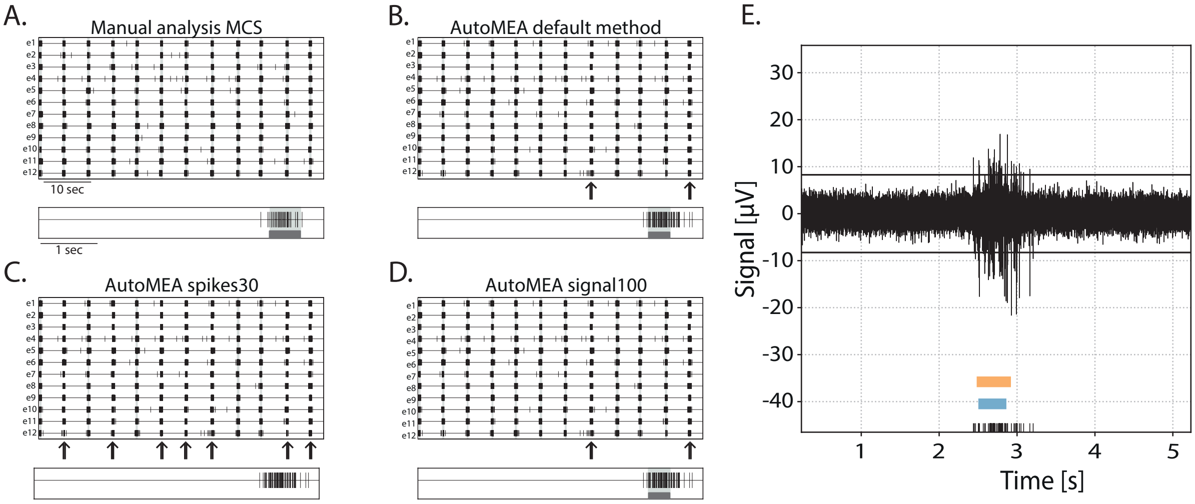


Figure 4: Example of outlier neuronal network in the cortical dataset A-D) Rasterplot and 5 second zoom in of a single electrode of spikes and bursts detected by manual analysis (A), autoMEA default method (B), spikes30 model (C), and signal100 model (D). Black arrows represent network bursts that are not detected in using the autoMEA software. E) raw trace of a burst that was not accurately detected using the autoMEA software. black lines at the bottom represent detected spikes, blue bar (bottom) represents burst as detected using the manual analysis, orange bar (top) represents burst as detected using the default method.
